# Presence and the global implications of plastics in wild commercial fish in the Alboran Sea

**DOI:** 10.1101/2021.12.11.472227

**Authors:** Sergio López-Martínez, Cipriano Perez-Rubín, Rafael Gavara, Rebecca Handcock, Marga L Rivas

## Abstract

The presence of plastic in the environment has become a major problem for marine megafauna. The identification of the global micro and mesoplastic uptake by commercial fish populations may allow for a better understanding of their impact. This study aims to: (i) determine the presence and composition of plastic in two pelagic fish (*Engraulis encrasicolus* and *Scomber scombrus*) and two demersal species (*Scyliorinus canicula* and *Mullus barbatus*) from the Alboran Sea (western Mediterranean) to quantify the relationship between plastic prevalence and habitat and feeding behavior in the selected fish species, (ii) compare local measurements made of the presence of plastics ingested by these four fish species with published values from a across their range literature review, and (iii) identify the methodologies used in similar studies of plastic pollution in fish. Across their range, the highest occurrence of plastics was found in *E. encrasicolus*, which contrasts to that found in *S. scombrus* at the Alboran sea. Material analysis of the collected data showed the most predominant fiber color was black and the predominant plastic polymer was polyethylene. The increasing emerging risk of plastics and the levels of macro- and micro-plastic ingested by seafood in this study support the suggest that quantifying plastic presence and composition may be essential as a food safety measure.

## 1. Introduction

Marine plastic pollution has increased tenfold since 1980, affecting the marine food web (Geyer et al., 2017). The combined and growing evidence of the magnitude and impacts of marine pollution from plastic waste (Bucci et al., 2020) is calling for urgent management strategies to address this problem (Bessa et al., 2019). Due to the increasing global synthetic fibers, microplastics (MPs) from synthetic textiles are likely to end up in the environment (Henry et al., 2019) and the ocean in particular (Almroth et al., 2018) mainly through washing processes and wastewater treatment plants (de Falco et al., 2019). Lima et al. (2021) estimated that globally there are approximately 5900□±□6800 m^−3^ microfibers in the ocean and recently it was confirmed that the highest concentrations of MPs are found on the seafloor (Kane et al., 2020). Such large plastic waste concentrations are not limited to areas of high population density but also occur in the most remote areas on the planet, from Antarctica and the Arctic Ocean (Lima et al., 2021), to abyssal depths (Peng et al., 2020). Recently, MPs ingestion by marine biota has gained more attention because of their increased bioavailability through fragmentation (Wesch et al., 2016) and consequently, the risks they may pose to the marine environment, such as leaching of additives (Koelmans et al., 2014). Thus, the ingestion and accumulation of plastics made from artificial polymers such as polyethylene (PE), polypropylene (PP), polystyrene (PS), polyvinyl chloride (PVC), polyurethane (PU) and polyethylene terephthalate (PET) (Pitt et al., 2018) also represents an indirect pathway of exposure to pollutants (Bellas and Gil, 2020) and might have negative impacts on animal health (Schrank et al., 2019).

A wide range of marine species have been documented to ingest plastics (Lopez-Martinez et al., 2021), with unpredictable consequences at the ecosystem level (Law et al., 2014; Beaumont et al., 2019). Commercial wild fish are key species in ecosystems and an essential and important source of food (FAO, 2016).

The first studies on plastic ingestion by wild marine fish were published in the early 1970s (Markic et al., 2020), but to date they were mainly focused on monitoring the uptake of meso- and macro-debris (Ajith et al. 2020; Foley et al. 2018; Barnes et al., 2009).

The ingestion of plastic microfibers by numerous commercial fish species has been recently studied (Rodriguez-Romeu et al., 2020), and even small MPs (< 3 mm) have been found in the tissues of commercial fish species, such as *Serranus scriba* (Zitouni et al., 2020). However, as far as we know, there is not yet a global overview of the extent of the threat of MPs to commercial marine fish species, mainly because the plastic uptake is dependent on the characteristics of each fish species and location (Provencher et al., 2019).

The identification of the variability of plastics uptake by different commercial marine fish species with different roles in the food web will allow for a better understanding of the impact that plastics have at the level of the individual, species, habitat, and food web. Hence, this study aims to: (i) determine the presence and composition of plastic in two pelagic fish (*Engraulis encrasicolus* and *Scomber scombrus*) and two demersal species (*Scyliorinus canicula* and *Mullus barbatus*) from the Alboran Sea (western Mediterranean) to quantify the relationship between plastic prevalence and habitat and feeding behavior of the selected fish species, (ii) compare local measurements made of the presence of plastics ingested by these four fish species with published values from across their range literature review, and (iii) identify the methodologies used in similar studies of plastic pollution in fish.

## 2. Methodology

### 2.1 Species and site data collection

Chondrichthyes and Osteichthyes species were selected for this study following the criteria for good species indicators as advised by Group of Experts on Scientific Aspects of Marine Environmental Protection (GESAMP, 2019). According to the categories of www.fishbase.se (Froese and Pauly, 2021), selected species should ideally be abundant, commercially and ecologically important, have a regional representation and a variety of feeding strategies, or several ecological niches like demersal and pelagic habitats. Our species-selection assessment covered 194 specimens of two pelagic species, *E. encrasicolus* (European anchovy) and *S. scombrus* (Atlantic mackerel), and two demersal species *S. canicula* (Lesser spotted dogfish) and *M. barbatus* (Red mullet). The specimens were obtained by commercial fishing boats caught in the northern Alboran Sea which is part of the Geographical Sub-Area 1 (GSA 1) identified by the Food and Agriculture Organization’s (FAO) General Fisheries Commission for the Mediterranean (GFCM), and in the west of the Western Mediterranean Sea sub-region according to the Marine Strategy Framework Directive (MSFD). In addition to their commercial importance, half of the species present pelagic ecological and habitat strategies and the other two are demersal. Fish samples were collected over a period of 4 months (October 2020 to January 2021) from local fisherman in an area comprised between the Gulf of Almeria (Table 1).

**Table 1.** Specimens collected by commercial fishing boats at the Alboran Sea (Western Mediterranean) from October 2020 to January 2021

| Common name | Species | N | Weight (g) | Size (cm) | Habitat | Location |  | Depth | % specimens with fibers | Total n of fibers |
| --- | --- | --- | --- | --- | --- | --- | --- | --- | --- | --- |
|  |  |  |  |  |  | Latitude | Longitude |  |  |  |
| European anchovy | *E. encrasicolus* | 51 | 27.60 ± 3.13 | 15.89 ± 0.64 | Pelagic | 36.772785 | -2.513525 | 40 | 0.0 | 0 |
| Atlantic mackerel | *S. scombrus* | 40 | 193.66 ± 24.34 | 27.12 ± 1.13 | Pelagic | 36.749861 | -2.549285 | 33 | 37.5 | 15 |
| Lesser spotted dogfish | *S. canicula* | 51 | 145.22 ± 40.65 | 37.29 ± 4.86 | Demersal | 36.46761 | -2.21380 | 85 | 9.8 | 7 |
| Red mullet | *M. barbatus* | 52 | 62.06 ± 20.58 | 17.71 ± 1.54 | Demersal | 36.46761 | -2.21380 | 85 | 32.7 | 24 |

### 2.2 Sample processing

In this study, fish captured for sample processing were stored at −20 °C until further analysis in the laboratory. During the laboratory analysis the stored captured fish were thawed for the extraction of the digestive tract, according to a previously published methodology (Lusher et al., 2013; Rocha-Santos and Duarte, 2015). The digestive tracts were stored in glass storage jars for subsequent identification of microplastics. Every specimen was measured and weighed (1 mm and 0.01 g of precision, respectively). The gastrointestinal tract was extracted and weighed, put in a glass container and digested with 10% potassium hydroxide (KOH) solution (Foekema et al., 2013) pre-filtered with a 150 μm sieve. The samples were placed in an oven for 48 hours at 40 °C (Bessa et al., 2019). Once the digestion was completed each sample was homogenized with a magnetic agitator.

To prevent and control airborne contamination during the sample processing procedure, all glassware, equipment and lab surfaces were cleaned with ethanol at 70% pre-filtered with a 150 μm sieve. For this study, all the sample processing was performed in a separate room of the laboratory, which was previously cleaned with 70 % ethanol and a cotton lab coat was used at all times. To identify possible airborne contamination, three glass microfiber filters placed in Petri dishes were opened every time the digestive tract or the digested mixture was exposed to air.

### 2.3 Physical analysis

The shape, size and color of plastic particles are important factors as they can affect the probability of their being encountered or ingested by marine organisms (e.g. de Sá *et al*., 2015). The visual identification of plastic particles is mainly done manually with a stereo microscope, sometimes with ultraviolet light or through a scanning electron microscope. In the present study, we used the most common categories in the GESAMP (2019) protocol to classify plastics by shape as fragments, foams, films, fibers/filaments, or pellets. In terms of size, the protocol recommends using the terms mega for >1 m, macro for 25 mm to 1 m, meso for 5 to 25 mm, and micro for 1 to 5 mm (GESAMP, 2019). The color of the particle may provide useful information and can potentially identify the preferred feeding strategies of some organisms, or the conditions to which the plastic particles have been exposed to. However, visual identification of color is subjective (GESAMP, 2019), and as there is currently no standard scheme for the color designation of plastic debris (Marti et al., 2020) in our analysis we identified common colors.

For our samples, the digested solution was poured into a sterilized glass Petri dish (20 cm diameter) with a cover to avoid airborne contamination. The identification of potential plastic particles was performed with a LEICA stereo microscope. Once potential plastic particles were identified, they were isolated and placed on a glass microfiber filter VWR®Glass Fiber Filters 1.3 μm pore size Each particle was characterized by color, type and size using a stereomicroscope with the camera LEICA MC 190 HD and the software ImageJ v.1.8.0_172 (Rasband W. S., 2011).

### 2.4 Chemical analysis

Chemical characterization of plastic samples is key to identifying potential plastic sources. The types of polymers that are commonly encountered in the marine environment are: low-density polyethylene (LDPE), high-density polyethylene (HDPE), PP, PS, PA, polyethylene terephthalate (PET), PVC, and cellulose acetate (CA) (Andrady, 2011). Renner et al. (2018) highlighted that the most commonly used analytical technique to identify the composition of macro- and micro-plastics in environmental samples is through FTIR or Raman spectroscopy. Both methods are very accurate in terms of polymer identification and have the advantage of being non-destructive (Prata et al., 2019). We therefore used μFTIR for polymer identification of the items found in our samples since all the plastics found were fibers and FTIR does not allow to identify fibers efficiently (Zhang et al., 2021). A Jasco model 4100 FTIR spectrophotometer (JASCO ANALITICA SPAIN, Madrid) equipped with a MIRacle Single Reflection ATR and a ZnSe crystal (Pike Technologies, Madison WI) was used for polymer identification of the items found in our samples. The resolution was 4 cm–1 in the range of 4000 to 600 cm–1 and 32 scans were recorded per sample.

### 2.5 Collating published studies of plastic in the four fish species in the study

To compare the presence of plastic in the four fish species that we analyzed from our study sites we collated published values from a literature collection that created from reviewing available literature. The aim of this literature review was to collate the data that had already been published rather than being a systematic review of the effects of plastic pollution such as by Bucci et al. (2020). The literature collection was generated by searching the global Web of Science, Scopus, and Google Scholar citation indexing databases for all existing publications up until April 2021. The keywords used for searching were combinations of each of our four fish species names of “*Engraulis encrasicolus*”, “*Scomber scombrus*”, “*Scyliorinus canicula*” and “*Mullus barbatus*”, with the additional search terms of “plastic”, “debris” and “plastic ingestion”.

To explore the characterization of plastics in our four fish species through the data from the literature collection, for each published study we first identified in each whether the stomach or the whole gastrointestinal tract was used in sample processing. Second, we identified the sample processing techniques used in the published study, categorized as visual observation, density separation, alkaline digestion and others (GESAMP, 2019), and the analysis method classified as visual identification, Fourier Transform Infrared (FTIR), microFTIR and Raman spectroscopy. The data from the published studies in the literature collection was also used to compile global data on the ingestion of macro- meso- and micro-plastics by these four fish species in ocean basins and seas.

### 2.6 Statistical analysis

Statistical analysis of the collated data used non-parametric methods after the data was found to not be normally distributed using Shapiro Wilk test. The Kruskal-Wallis test for multiple comparisons was used with a significance level of 0.05. A Chi square test was used to find differences among species, colors and polymer types, and data from the literature collection. A U Mann-Whitney test was used to compare differences between groups. A Generalized Linear Model (GLM) was used to estimate size effects. Statistical analysis was performed using the R studio version 4.0.2 (2020-06-22) (2020) software.

## 3. Results

### 3.1 Physical and chemical characterization

Our study data showed that from the total of 194 specimens sampled from October 2020 to January 2021 at the Alboran Sea, *S. scombrus* showed the highest plastic ingestion with 37.5 % of specimens containing synthetic fibers, while none were found in *E. encrasicolus* (Fig 1, Table 1). A total of 19 fibers smaller than 1 mm were not measured because of similarities with identified airborne contamination. By contrast, the average size of items larger than 1 mm was 4.56 ± 2.68 mm (n = 36), and no significant differences between fiber size among species was found (X = 64, p = 0.27). Black was the most predominant color (46.3 %, n = 54), followed by blue (Fig 2i (f)). However, for the polymer predominance, cellulose was the most frequently identified, followed by PET polymer (Fig. 2ii (f), Table 1 SI).

**Figure 1.**
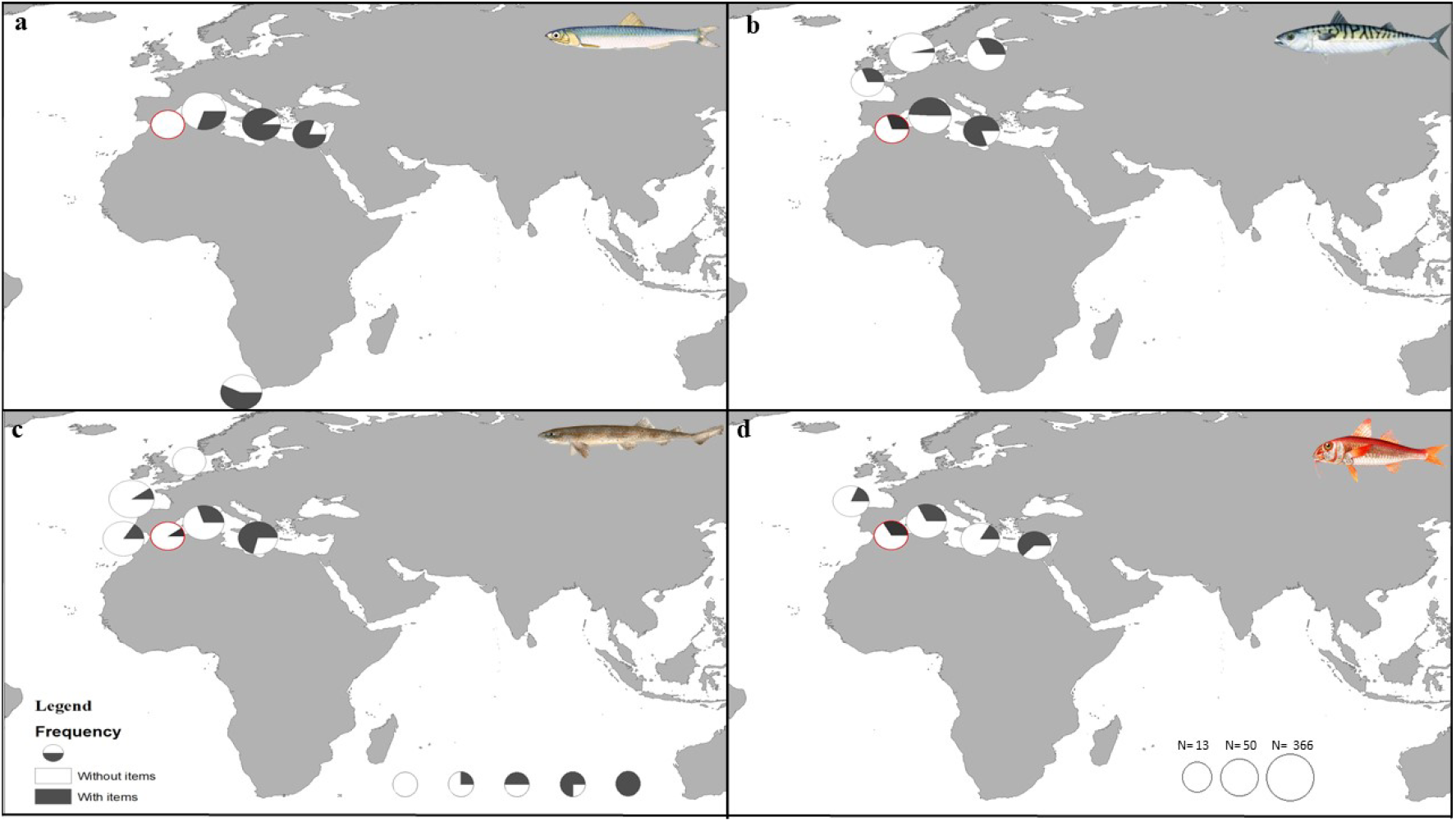
Frequency of ingested plastic in (a) *E. encrasicolus*, (b) *S. scombrus*, (c) *S. canicula* and (d) *M. barbatus*. In each circle, the black portion represents the percentage of plastic found in that area. The size of the circle is proportional to the number of individuals. The red circle shows the Alboran Sea study location.

**Figure 2.**
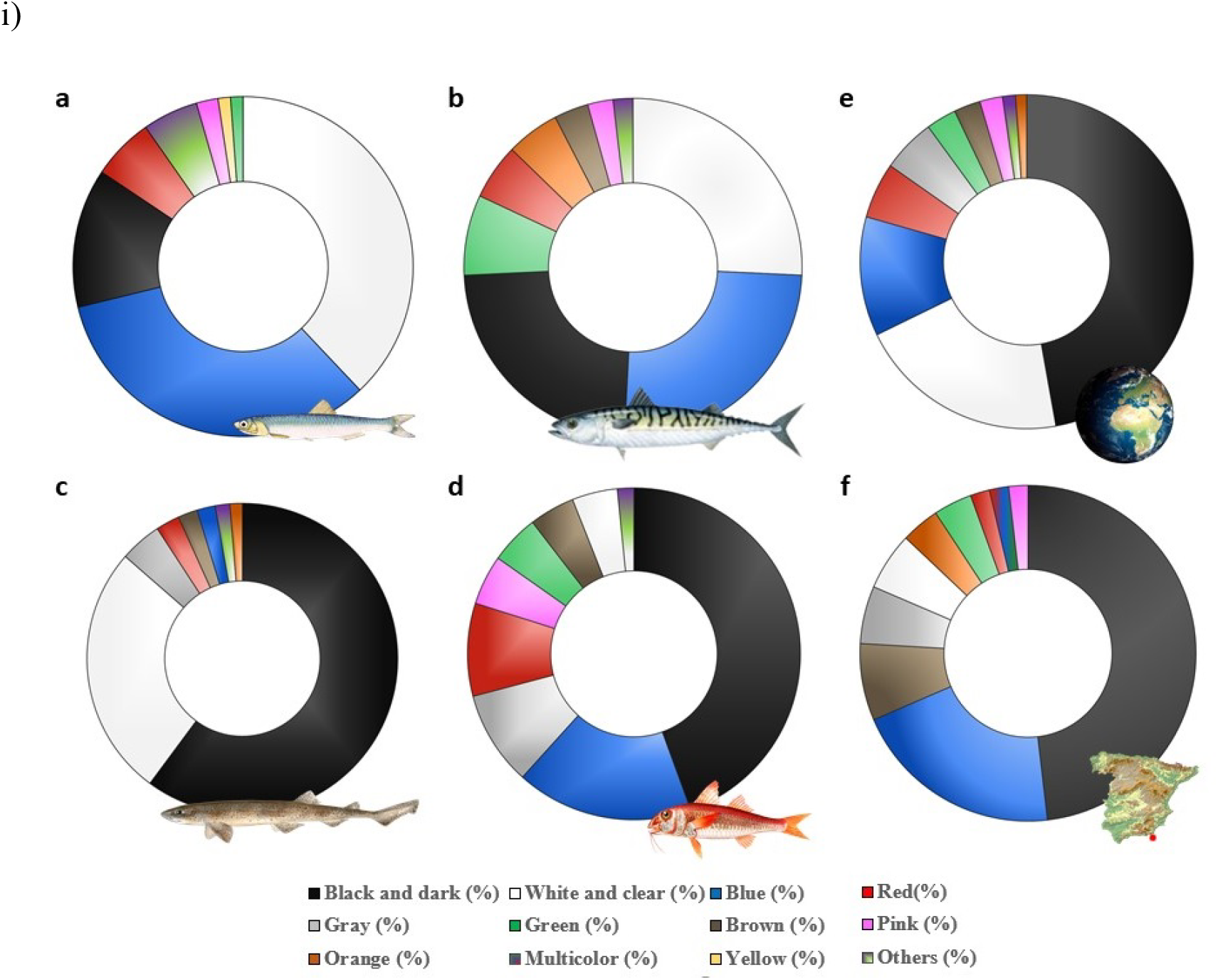
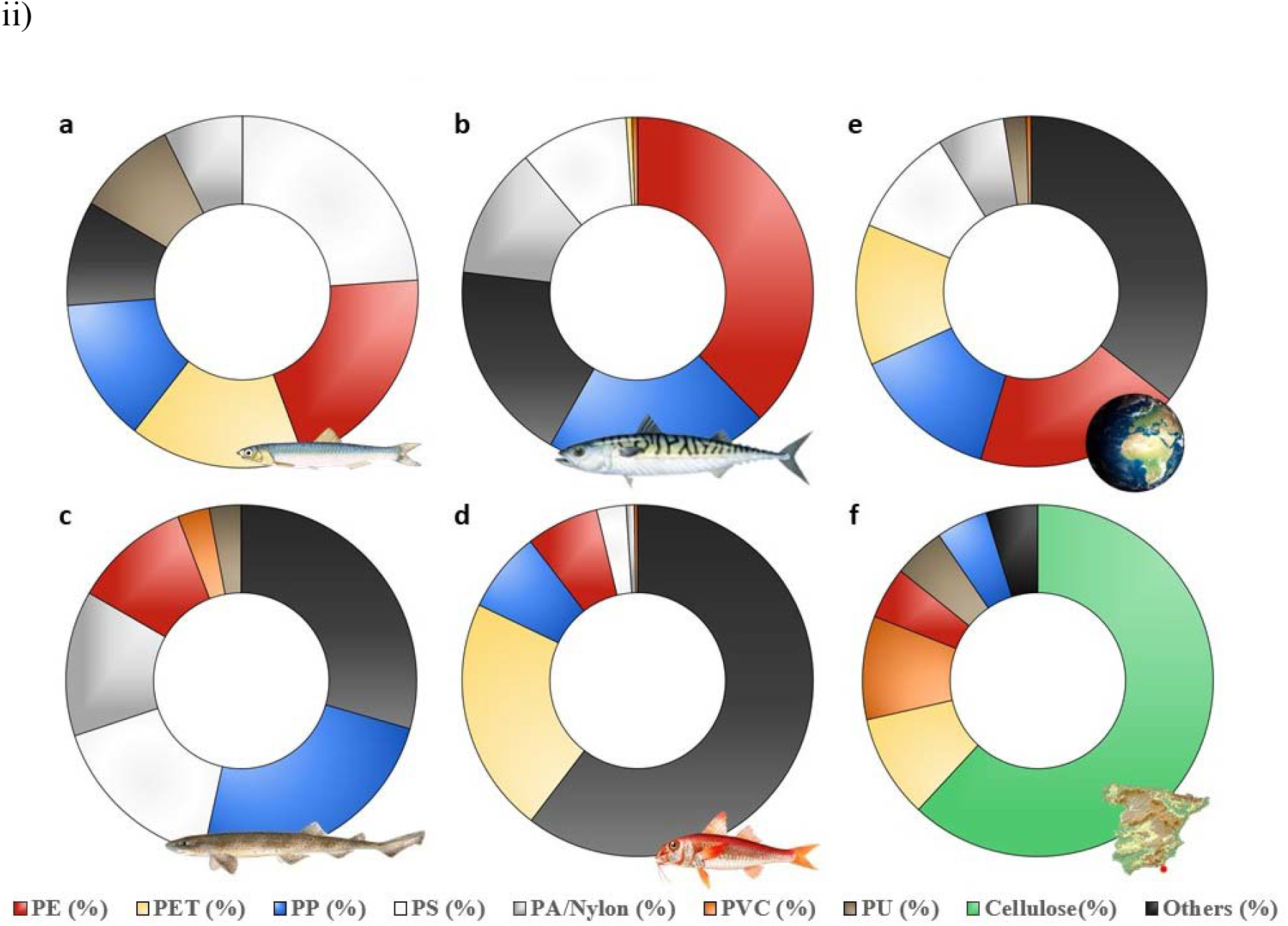
Radial plots of (i) fiber color by species: (a) *E. encrasicolus*, (b) *S. scombrus*, (c) *S. canicula*, and (d) *M. barbatus* at the total global scale and (e) color across their range and (f) local scales, and (ii) polymer composition by species: (a) *E. encrasicolus*, (b) *S. scombrus*, (c) *S. canicula* and (d) *M. barbatus* at the total global scale, and (e) color across their range and (f) local scale.

Airborne contamination was detected in the blanks, all of which were fibers, with 70.3 % (n =57) being smaller than 1 mm. 4 fibers larger than 1 mm were detected in the *E. encrasicolus* control blanks, 16 fibers in *S. scombrus*, 4 fibers in *S. canicula* and none in *M. barbatus*. Most of these fibers were of dark colors.

Our investigation into whether the presence of plastics was dependent on whether the species were pelagic or demersal found no significant differences for plastic uptake between both groups (X = 2.25, p = 0.13).

### 3.2 Comparing observations with studies from the collated literature

A total of 36 candidate publications about the selected studied species were identified from the citation indexing databases as of April 2021. Specifically, there were 12 studies for *E. encrasicolus*, 5 studies for *S. scombrus*, 10 studies for *S. canicula*, and 9 studies for *M. barbatus*.

Across their distribution range, the highest occurrence of plastic abundance was found in *E. encrasicolus* with 42.08 ± 30.75 % (average ± SD) (n = 766) of specimens with plastics (Fig 1a), showing significant differences among species (X = 174.1, p < 0.01) (Fig. 2a).

Most of the plastics found across the range of distribution in the selected species were fibers. Significant differences were found in the color of the fibers ingested by fish. Differences were found among species (X = 364.9, p < 0.01) (Fig. 3a) and the presence of plastics was higher in pelagic than demersal groups (U Mann-Whitney test = 622148, p < 0.01) (Fig. 3b). Across their range and by species, the predominant fiber color was black (Fig. 2i (e)). The most frequent polymer identified by FTIR was PE (22.42 %) (Fig 2ii (e)). No significant differences were found for polymer composition among species (X = 248.2, p < 0.01) and groups (X = 37.1, p < 0.01).

**Figure 3.**
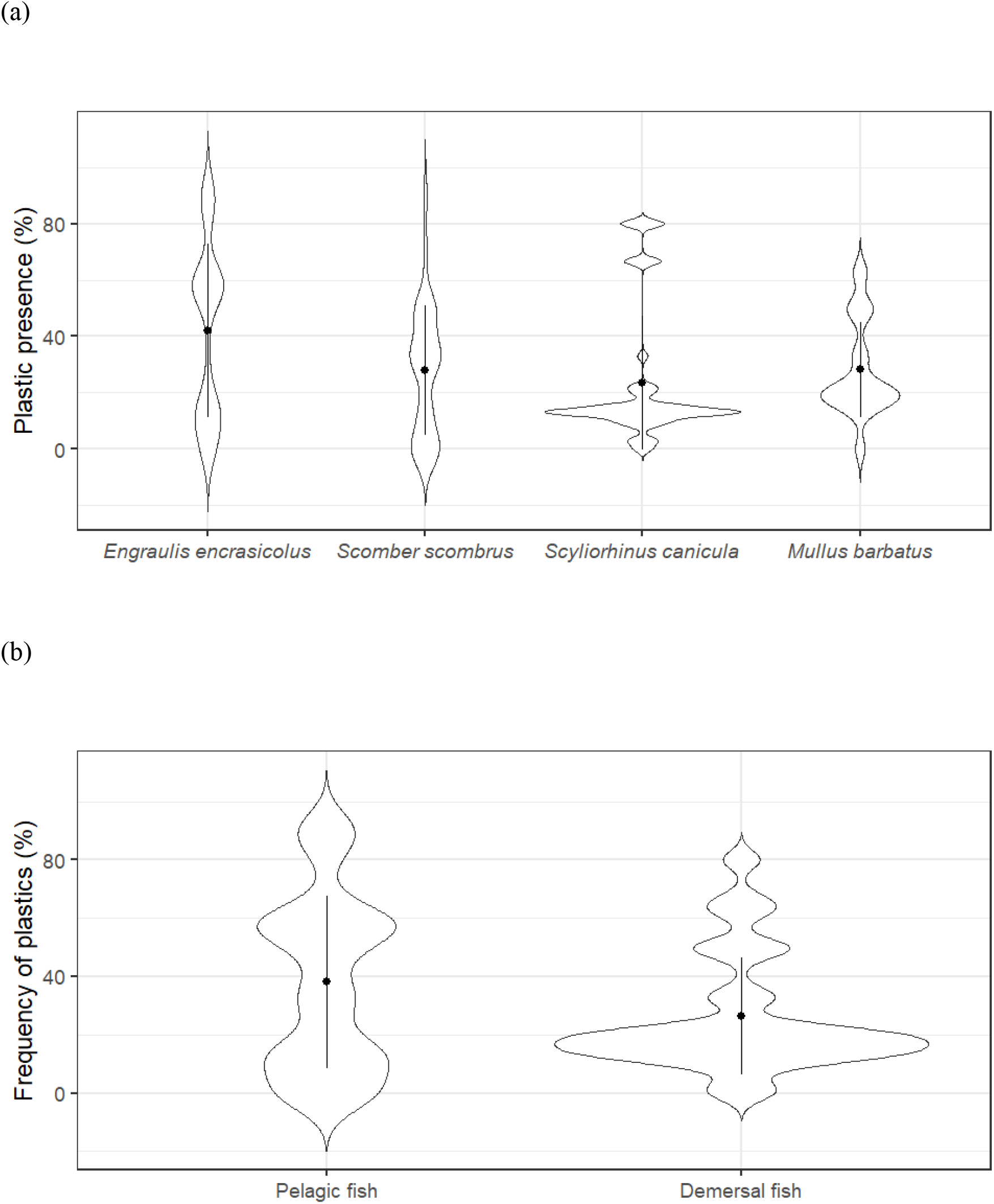
Violin plots of frequency (%) of plastic presence by (a) species and (b) habitat across their range. The black point is the average of plastic frequency, the black lines indicate the standard deviation, and the width of the violin represents the sample size.

**Figure 4.**
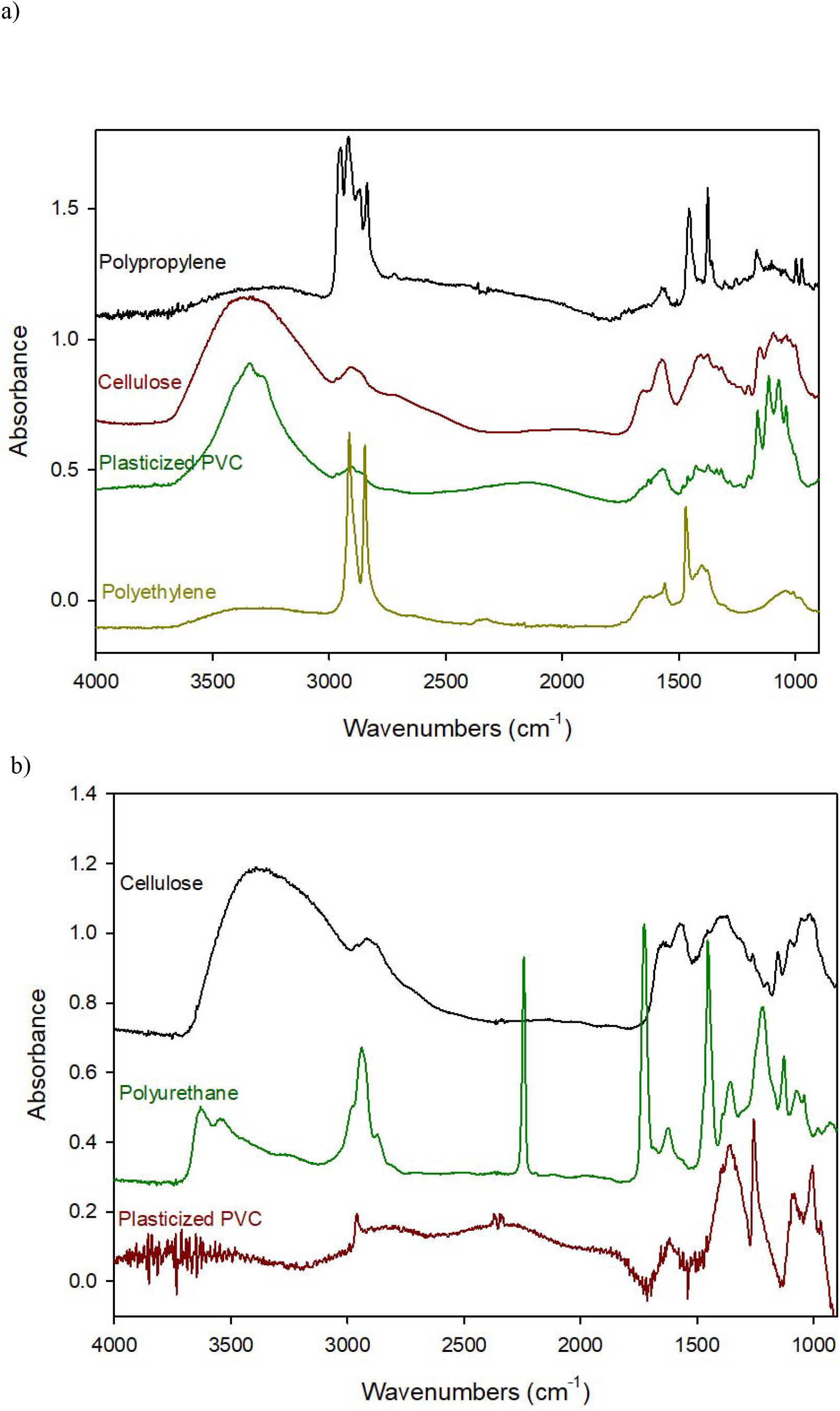
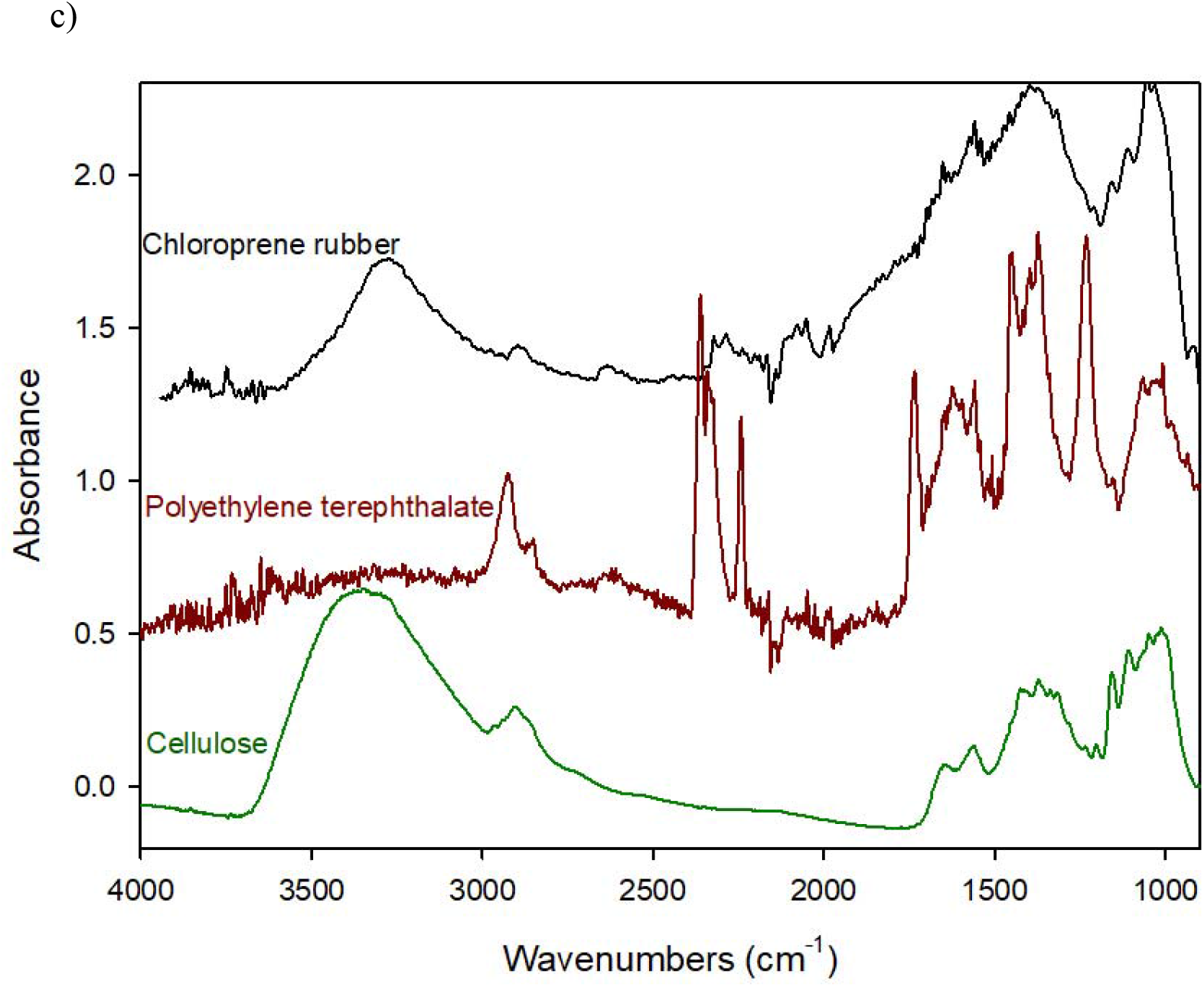
FTIR-ATR spectra of microplastics found in samples (a) *S. scombrus*, (b) *S. canicula* and (c) *M. barbatus* and their identification. Spectra have been shifted vertically to provide a better view of the different signals.

Our data in the Alboran Sea showed significant differences in the presence of plastic which increased for the Mediterranean locations to the east (X = 13.66, p = 0.003). In addition, the presence of plastics in pelagic or demersal groups differed significantly between our results and over their range of distribution (p = 0.07).

There were no differences between the type and color of plastic ingestion found in our results and those found in the literature, where all plastics found were also fibers and black was also the most frequent color (X = 10.05, p = 0.35) (Fig 2i (e, f)), except to the polymer composition where the Alboran Sea showed that highest presence of cellulose rather than PE (X = 409.5, p < 0.01)(Fig 2ii (e, f)).

### 3.3 The methodologies used in plastic pollution research

In the total set of candidate publications in the literature collection, 12 studies dissected the stomach of the fish for physical analysis, and 24 the digestive tract. The most used chemical analysis technique was FTIR (including ATR and μFTIR) with a total of 17 studies, followed by visual analysis and Raman spectroscopy, with 8 and 7 studies respectively (Table 1 SI).

Overall, the small size of the literature collection highlights that it is still a great challenge to find scholarly publications with a standardized methodology for the detection of plastic ingestion in fish. For instance, it was not possible to estimate overall averages of plastic size across the dataset because some studies included size ranges (maximum range 0.001 - 19 mm) and others just the size averages. Additionally, a high percentage of polymers were included as “others”, which could be due to the absence of chemical analysis in those studies or to the difficulties encountered in the identification of polymers by the technique used.

It has also been noted that there was large variability in the number of ingested plastic items depending on the sampling site, and possibly also influenced by the time of day of capture (Table 1 SI).

Finally, it should be highlighted the absence of control airborne contamination measurements with glass microfibers in most of the studies, which it could have overestimated results (Table 1 SI).

## 4. Discussion

In this study we identified and characterized the plastic content in four wild commercial fish from a Mediterranean area, the Alboran Sea and correlated this with published values across their range based on a literature collection of published studies.

Our results across the fish distribution range showed that the studied species in the Alboran Sea have relatively medium concentrations of plastics, and an increase in the frequency of plastic presence increases longitudinally when moving west towards the eastern Mediterranean Sea. The Mediterranean Sea does have low water circulation and is surrounded by such a high population density, making it one of the most polluted seas on Earth (López-Martínez et al., 2020). The spatial distribution of MPs in the global seas and oceans has been recognized is concentrated in areas of convergence and accumulation such as gyres (Cózar et al., 2015), as well as enclosed bays, gulfs and seas surrounded by densely populated coastlines as in the case of the Mediterranean Sea (Collignon et al., 2012; Reisser et al., 2013; Palazzo et al. 2021). This hydrodynamic pattern of plastics suggests that a part of the floating plastic pollution in the Mediterranean may originate outside the basin, with the Mediterranean Sea acting as a sink for floating plastic pollution from the Atlantic (Béranger, K et al. 2010; Aliani et al. 2003). Models from Cózar et al. in (2015) and Eriksen, et al. in (2014) showed a gradual increase of the number plastic parts from west to east of the Mediterranean. However, as reported by Cózar et al. (2015), broad-scale plastic distribution models should be treated with caution due to the low agreement found between measurements and scale-dependent model predictions. In our study, we validated these models of plastic quantity in biota as proxies for plastic quantity in habitat. The development of more accurate estimates on the magnitude and distribution of plastic pollution in the Mediterranean Sea requires increased spatial and temporal resolution and coverage.

It is expected that due to the density characteristics of MPs, they would accumulate on the seafloor where conditions allow, becoming a sink for plastic particles (Woodall et al., 2014). However, in the water column MPs are distributed according to various intrinsic factors such as particle density, size and shape (Ballent et al., 2012; Enders et al., 2015) and extrinsic factors such as wave movements, currents and storms (Kukulka et al., 2012; Reisser et al., 2015). The physical properties of synthetic fibers allow them to remain in the water’s upper layers for longer (Ballent et al., 2012; Reisser et al., 2013). This may be the reason why the particles found in pelagic species in this study might have high bioavailability, and consequently high ingestion. Fibers were the predominant MPs type detected in our samples, resembling what has been found in other studies on MPs ingestion in fish (Avio et al., 2015; Bellas et al., 2016; Peters et al., 2017; Ory et al., 2018; Compa et al., 2018; Ugwu et al., 2021, Palazzo et al., 2021). Although in our sampling, *E. encrasicolus* did not contain fibers, the assessment of the 4 studied species across their range found this was the one with the most prevalent fibers ingested. Pelagic planktivorous species like fish are susceptible to accumulate MPs, since their prey have the same size range and MPs tend to accumulate in the same feeding areas and spawning grounds (Lopes et al., 2020). Therefore, it is essential to correlate plastic ingestion with habitat and fish’s feeding behavior, water column distribution and geographic areas to be able to map in detail how plastic might be interacting with species and where it might accumulate in the long term.

We also found a wide disparity of plastic sizes in the studies, with our data falling within the common size range found in other publications. We found that black was the predominant color of plastics in our data, as was also found in other studies (Capillo et al. 2020, Savoca et al. 2020, Bellas et al., 2016), Probably, part of the ingestion of remains occurs unintentionally (Lusher et al. 2013), or as suggested by Boerger et al., (2010), which could be due to their feeding behaviors, in which they could confuse the color of the fibers with that of the prey.

The results across the ranges of our sampled species on the origin of plastics point to PE as the most commonly polymer ingested by marine vertebrates (Lopez-Martinez et al., 2020, Ugwu et al., 2021). In contrast, our results showed that cellulose was the most predominant polymer in the Alboran Sea, which may have been because it was included as ‘others’ in many of the other studies. The noise level of the results varied due to different sample sizes. In some cases, the sample was very narrow and produced higher noise levels. However, since the use of FTIR was for identification purposes, the presence of noise was not relevant to the interpretation of the spectra. PE absorbs more hydrophobic chemicals, such as polychlorinated biphenyls (PCBs), than other polymers (Teuten et al., 2009). Additionally, MPs contain additives and hazardous chemicals that are added during plastics manufacture, while other MPs can adsorb it from the water (Teuten et al., 2009). Therefore, the consumption of fish contaminated with MPs may lead to a higher probability of exposure to bisphenols and other endocrin disruptors (Barboza et al 2020). It is therefore essential to know the composition of each polymer’s additives because this varies among different types of plastic. Levels of liver oxidative stress and inflammation and in internal tissues have also been correlated with higher microplastic intake (Capó et al., 2021). Smaller particles such as nanoplastics have the potential to affect the composition, diversity and functionality of the gut microbiome of vertebrate and invertebrate organisms. When the composition of the gut microbiome is modified in situations of repeated and persistent exposure to nano-plastics, alterations arise in the immune, endocrine and nervous systems (Teles et al., 2020).

## 5. Limitations and recommendations

We identified a small set publications with a standardized methodology for the detection of microplastics in fish as Bessa reported in (2019), which makes comparisons difficult among individual sites and for global assessments. Considering that most studies did not include chemical analysis and the difficulties that were encountered in identifying polymers with the FTIR technique, we recommend using chemical analysis with techniques such as FTIR or Raman spectroscopy, which gives accurate results and are recommended by experts (GESAMP).

One limitation of this study is the large variability in the observed number of ingested plastic items that was found among sampling site, which may also have been dependent on the time of day of capture. Thus, in any future studies the methodological sampling of bioindicator species should also be further developed.

Our assessment of the collated research showed that although the 4 species in this study have a wide distribution, there is a lack of studies from Low-Income Countries and most of studies were focused on the Mediterranean Sea and the North Atlantic. We therefore recommend the continued monitoring of the plastic prevalence in areas such as Africa and remote areas where the presence of the plastics in the study species has still not been assessed. This knowledge gap on plastic ingestion by the study species across their ranges also suggests careful use of global assessments of plastic ingestion.

This data on four fish species collected in this study adds to the body of research on plastics in wild commercial fish. The reviewed literature is necessarily focused on these species while complementing broader reviews of plastic pollution in marine, freshwater, and terrestrial environments (Bucci et al., 2020). The range of plastic characteristics that we measured such as shape, color, etc. also meets the call for more complex studies of microplastics (Bucci et al., 2020).

The species selected in this study are representative of two different feeding strategies that can potentially give us a better understanding of the distribution, characterization, fate and accumulation of MPs (Fossi et al., 2018). Preventive measures to minimize the release of these emerging pollutants into the oceans must be of concern and a conservation priority for every country. For instance, the implementation of filters with pore size of 60□μm in washing machines could retain and decrease the numbers of synthetic fibers released, which might significantly reduce them from wastewater releases into the oceans (de Falco et al 2019).

## Acknowledgments

This study has been carried out for the PlastiMarMed project supported by Fundación Biodiversidad through the Spanish Ministry for the Ecological Transition and Demographic Challenge (MITECO) and a FPU grant by the Spanish Ministry of Education, Innovation and Universities (aid for university teacher training [FPU19/ 05660]. We also give thanks to Francisca Ribeiro from the University of Queensland and Exeter, for his advice and review, Luis Hidalgo Oller and Jorge Herrera-González for specimens support, and Dr. Irene Rodriguez Dominguez from University of Almería for chemical technical advice.

## References

Ajith, N., Arumugam, S., Parthasarathy, S., Manupoori, S., & Janakiraman, S. 2020. Global distribution of microplastics and its impact on marine environment review. Environmental Science and Pollution Research, 27(21), 25970–25986.

Aliani, S., Griffa, A., & Molcard, A. 2003) Floating debris in the Ligurian Sea, north-western Mediterranean. Marine Pollution Bulletin, 46(9), 1142–1149.

Andrady, A.L., 2011. Microplastics in the marine environment. Mar. Pollut. Bull. 62, 1596–1605. https://doi.org/10.1016/j.marpolbul.2011.05.030

Ballent, A., Purser, A., Mendes, P.D.J., Pando, S., Thomsen, L., 2012. Physical transport properties of marine microplastic pollution. Biogeosciences Discuss. 9, 18755–18798. https://doi.org/10.5194/bgd-9-18755-2012

Barboza, L.G.A., Cunha, S.C., Monteiro, C., Fernandes, J.O., Guilhermino, L., 2020. Bisphenol A and its analogs in muscle and liver of fish from the North East Atlantic Ocean in relation to microplastic contamination. Exposure and risk to human consumers. J. Hazard. Mater. 393, 122419. https://doi.org/10.1016/j.jhazmat.2020.122419

Barnes, D.K.A., Galgani, F., Thompson, R.C., Barlaz, M., 2009. Accumulation and fragmentation of plastic debris in global environments. Philos. Trans. R. Soc. B Biol. Sci. 364, 1985–1998. https://doi.org/10.1098/rstb.2008.0205

Beaumont, N.J., Aanesen, M., Austen, M.C., Börger, T., Clark, J.R., Cole, M., Hooper, T., Lindeque, P.K., Pascoe, C., Wyles, K.J., 2019. Global ecological, social and economic impacts of marine plastic. Mar. Pollut. Bull. 142, 189–195. https://doi.org/10.1016/j.marpolbul.2019.03.022

Bellas, J., Gil, I., 2020. Polyethylene microplastics increase the toxicity of chlorpyrifos to the marine copepod Acartia tonsa. Environ. Pollut. 260, 114059. https://doi.org/10.1016/j.envpol.2020.114059

Bellas, J., Martínez-Armental, J., Martínez-Cámara, A., Besada, V., Martínez-Gómez, C., 2016. Ingestion of microplastics by demersal fish from the Spanish Atlantic and Mediterranean coasts. Mar. Pollut. Bull. 109, 55–60. https://doi.org/10.1016/j.marpolbul.2016.06.026

Béranger, K., Drillet, Y., Houssais, M. N., Testor, P., Bourdallé-Badie, R., Alhammoud, B.,… & Crépon, M. (2010). Impact of the spatial distribution of the atmospheric forcing on water mass formation in the Mediterranean Sea. Journal of Geophysical Research: Oceans, 115(C12). https://doi.org/10.1029/2009JC005648

Bessa, F., 2019. Harmonized protocol for monitoring microplastics in biota Micropoll-Multilevel assessment of microplastics and associated pollutants in the Baltic Sea View project. https://doi.org/10.13140/RG.2.2.28588.72321/1

Boerger, C. M., Lattin, G. L., Moore, S. L., & Moore, C. J. (2010). Plastic ingestion by planktivorous fishes in the North Pacific Central Gyre. Marine pollution bulletin, 60(12), 2275–2278. https://doi.org/10.1016/j.marpolbul.2010.08.007

Bucci, K., M. Tulio, and C.M. Rochman. 2020. What is known and unknown about the effects of plastic pollution: A meta-analysis and systematic review. Ecological Applications 30(2):e02044, https://doi.org/10.1002/eap.2044.

Capillo, G., Savoca, S., Panarello, G., Mancuso, M., Branca, C., Romano, V.,… & Spano, N. 2020. Quali-quantitative analysis of plastics and synthetic microfibers found in demersal species from Southern Tyrrhenian Sea (Central Mediterranean). Marine pollution bulletin, 150, 110596. https://doi.org/10.1016/j.marpolbul.2019.110596

Capó, X., Company, J. J., Alomar, C., Compa, M., Sureda, A., Grau, A.,… & Deudero, S. (2021). Long-term exposure to virgin and seawater exposed microplastic enriched-diet causes liver oxidative stress and inflammation in gilthead seabream *Sparus aurata*, Linnaeus 1758. Sci. of The Total Environ. 767, 144976. https://doi.org/10.1016/j.scitotenv.2021.144976

Carney Almroth, B.M., Åström, L., Roslund, S., Petersson, H., Johansson, M., Persson, N.K., 2018. Quantifying shedding of synthetic fibers from textiles; a source of microplastics released into the environment. Environ. Sci. Pollut. Res. 25, 1191–1199. https://doi.org/10.1007/s11356-017-0528-7

Collignon, A., Hecq, J.H., Glagani, F., Voisin, P., Collard, F., Goffart, A., 2012. Neustonic microplastic and zooplankton in the North Western Mediterranean Sea. Mar. Pollut. Bull. 64, 861–864. https://doi.org/10.1016/j.marpolbul.2012.01.011

Compa, M., Ventero, A., Iglesias, M., Deudero, S., 2018. Ingestion of microplastics and natural fibers in Sardina pilchardus (Walbaum, 1792) and Engraulis encrasicolus (Linnaeus, 1758) along the Spanish Mediterranean coast. Mar. Pollut. Bull. 128, 89–96. https://doi.org/10.1016/j.marpolbul.2018.01.009

Cózar, A., Sanz-Martín, M., Martí, E., González-Gordillo, J. I., Ubeda, B., Gálvez, J. Á.,… & Duarte, C. M. 2015. Plastic accumulation in the Mediterranean Sea. PloS one, 10(4), e0121762. https://doi.org/10.1371/journal.pone.0121762

De Falco, F., Di Pace, E., Cocca, M., & Avella, M. 2019. The contribution of washing processes of synthetic clothes to microplastic pollution. Scientific reports, 9(1), 1–11. https://doi.org/10.1038/s41598-019-43023-x

de Sá, L.C., Luís, L.G., Guilhermino, L., 2015. Effects of microplastics on juveniles of the common goby (Pomatoschistus microps): Confusion with prey, reduction of the predatory performance and efficiency, and possible influence of developmental conditions. Environ. Pollut. 196, 359–362. https://doi.org/10.1016/j.envpol.2014.10.026

Enders, K., Lenz, R., Stedmon, C.A., Nielsen, T.G., 2015. Abundance, size and polymer composition of marine microplastics ≥10 μm in the Atlantic Ocean and their modelled vertical distribution. Mar. Pollut. Bull. 100, 70–81. https://doi.org/10.1016/j.marpolbul.2015.09.027

Eriksen, M., Lebreton, L. C., Carson, H. S., Thiel, M., Moore, C. J., Borerro, J. C.,… & Reisser, J. (2014). Plastic pollution in the world’s oceans: more than 5 trillion plastic pieces weighing over 250,000 tons afloat at sea. PloS one, 9(12), e111913.)

FAO. (2016). The State of World Fisheries and Aquaculture 2016 (SOFIA): Contributing to food security and nutrition for all. Rome. 200 pp.

Foekema, E.M., De Gruijter, C., Mergia, M.T., Van Franeker, J.A., Murk, A.J., Koelmans, A.A., 2013. Plastic in north sea fish. Environ. Sci. Technol. 47, 8818–8824. https://doi.org/10.1021/es400931b

Foley, C. J., Feiner, Z. S., Malinich, T. D., & Höök, T. O. 2018. A meta-analysis of the effects of exposure to microplastics on fish and aquatic invertebrates. Science of the total environment, 631, 550–559. https://doi.org/10.1016/j.scitotenv.2018.03.046

Fossi, M.C., Pedà, C., Compa, M., Tsangaris, C., Alomar, C., Claro, F., Ioakeimidis, C., Galgani, F., Hema, T., Deudero, S., Romeo, T., Battaglia, P., Andaloro, F., Caliani, I., Casini, S., Panti, C., Baini, M., 2018. Bioindicators for monitoring marine litter ingestion and its impacts on Mediterranean biodiversity. Environ. Pollut. 237, 1023–1040. https://doi.org/10.1016/j.envpol.2017.11.019

Froese, R., and D. Pauly. Editors. (2021). FishBase. World Wide Web electronic publication. www.fishbase.org, version (Accessed February 2021).

GESAMP, 2019. Guidelines for the monitoring and assessment of plastic litter in the ocean (Kershaw P.J., Turra A. and Galgani F. editors), (IMO/FAO/UNESCO-IOC/UNIDO/WMO/IAEA/UN/UNEP/UNDP/ISA Joint Group of Experts on the Scientific Aspects of Marine Environmental Prote. Rep. Stud. GESAMP no 99, 130p.

Geyer, R., Jambeck, J.R., Law, K.L., 2017. Production, use, and fate of all plastics ever made. Sci. Adv. 3, 25–29. https://doi.org/10.1126/sciadv.1700782

Henry, B., Laitala, K., Klepp, I.G., 2019. Microfibers from apparel and home textiles: Prospects for including microplastics in environmental sustainability assessment. Sci. Total Environ. 652, 483–494. https://doi.org/10.1016/j.scitotenv.2018.10.166

Kane, I.A., Clare, M.A., Miramontes, E., Wogelius, R., Rothwell, J.J., Garreau, P., Pohl, F., 2020. Seafloor microplastic hotspots controlle by deep-sea circulation. Science (80-.). 368, 1140–1145. https://doi.org/10.1126/science.aba5899

Koelmans, A.A., Besseling, E., Foekema, E.M., 2014. Leaching of plastic additives to marine organisms. Environ. Pollut. 187, 49–54. https://doi.org/10.1016/j.envpol.2013.12.013

Kukulka, T., Proskurowski, G., Morét-Ferguson, S., Meyer, D.W., Law, K.L., 2012. The effect of wind mixing on the vertical distribution of buoyant plastic debris. Geophys. Res. Lett. 39. https://doi.org/10.1029/2012GL051116

Law, K. L., & Thompson, R. C. 2014. Microplastics in the seas. Science, 345(6193), 144–145.https://doi.org/10.1126/science.1254065

Lima, A.R.A., Ferreira, G.V.B., Barrows, A.P.W., Christiansen, K.S., Treinish, G., Toshack, M.C., 2021. Global patterns for the spatial distribution of floating microfibers: Arctic Ocean as a potential accumulation zone. J. Hazard. Mater. 403. https://doi.org/10.1016/j.jhazmat.2020.123796

Lopes, C., Raimundo, J., Caetano, M., Garrido, S., 2020. Microplastic ingestion and diet composition of planktivorous fish. Limnol. Oceanogr. Lett. 5, 103–112. https://doi.org/10.1002/1012.10144

López-Martínez, S., Morales-Caselles, C., Kadar, J., Rivas, M.L., 2021. Overview of global status of plastic presence in marine vertebrates. Glob. Chang. Biol. 27, 728–737. https://doi.org/10.1111/gcb.15416

Lusher, A.L., McHugh, M., Thompson, R.C., 2013. Occurrence of microplastics in the gastrointestinal tract of pelagic and demersal fish from the English Channel. Mar. Pollut. Bull. 67, 94–99. https://doi.org/10.1016/j.marpolbul.2012.11.028

Markic, A., Gaertner, J.-C., Gaertner-Mazouni, N., Koelmans, A., 2019. Plastic ingestion by marine fish in the wild. Crit. Rev. Environ. Sci. Technol. 50, 1–41. https://doi.org/10.1080/10643389.2019.1631990

Martí, E., Martin, C., Galli, M., Echevarría, F., Duarte, C.M., Cózar, A., 2020. The Colors of the Ocean Plastics. Environ. Sci. Technol. 54, 6594–6601. https://doi.org/10.1021/acs.est.9b06400

Ory, N.C., Gallardo, C., Lenz, M., Thiel, M., 2018. Capture, swallowing, and egestion of microplastics by a planktivorous juvenile fish. Environ. Pollut. 240, 566–573. https://doi.org/10.1016/j.envpol.2018.04.093

Palazzo, L., Coppa, S., Camedda, A., Cocca, M., De Falco, F., Vianello, A., Massaro, G., de Lucia, G.A., 2021. A novel approach based on multiple fish species and water column compartments in assessing vertical microlitter distribution and composition. Environ. Pollut. 272, 116419. https://doi.org/10.1016/j.envpol.2020.116419

Peng, G., Bellerby, R., Zhang, F., Sun, X., Li, D., 2020. The ocean’s ultimate trashcan: Hadal trenches as major depositories for plastic pollution. Water Res. 168, 115121. https://doi.org/10.1016/j.watres.2019.115121

Peters, C.A., Thomas, P.A., Rieper, K.B., Bratton, S.P., 2017. Foraging preferences influence microplastic ingestion by six marine fish species from the Texas Gulf Coast. Mar. Pollut. Bull. 124, 82–88. https://doi.org/10.1016/j.marpolbul.2017.06.080

Pitt, J.A., Trevisan, R., Massarsky, A., Kozal, J.S., Levin, E.D., Di Giulio, R.T., 2018. Maternal transfer of nanoplastics to offspring in zebrafish (Danio rerio): A case study with nanopolystyrene. Sci. Total Environ. 643, 324–334. https://doi.org/10.1016/j.scitotenv.2018.06.186

Prata, J.C., da Costa, J.P., Girão, A. V., Lopes, I., Duarte, A.C., Rocha-Santos, T., 2019. Identifying a quick and efficient method of removing organic matter without damaging microplastic samples. Sci. Total Environ. 686, 131–139. https://doi.org/10.1016/j.scitotenv.2019.05.456

Provencher, J.F., Ammendolia, J., Rochman, C.M., Mallory, M.L., 2019. Assessing plastic debris in aquatic food webs: What we know and don’t know about uptake and trophic transfer. Environ. Rev. 27, 304–317. https://doi.org/10.1139/er-2018-0079

Rasband, W.S., 2018. ImageJ, U. S. National Institutes of Health, Bethesda, Maryland, USA, https://imagej.nih.gov/ij/, 1997–2018.

R Core Team 2020. R: A language and environment for statistical computing. R Foundation for Statistical Computing, Vienna, Austria. URL https://www.R-project.org/.

Reisser, J., Shaw, J., Wilcox, C., Hardesty, B.D., Proietti, M., Thums, M., Pattiaratchi, C., 2013. Marine Plastic Pollution in Waters around Australia: Characteristics, Concentrations, and Pathways. PLoS One 8, e80466.

Reisser, J., Slat, B., Noble, K., Du Plessis, K., Epp, M., Proietti, M., De Sonneville, J., Becker, T., Pattiaratchi, C., 2015. The vertical distribution of buoyant plastics at sea: An observational study in the North Atlantic Gyre. Biogeosciences 12, 1249–1256.https://doi.org/10.5194/bg-12-1249-2015

Renner, G., Schmidt, T.C., Schram, J., 2018. Analytical methodologies for monitoring micro(nano)plastics: Which are fit for purpose? Curr. Opin. Environ. Sci. Heal. 1, 55–61. https://doi.org/10.1016/j.coesh.2017.11.001

Rocha-Santos, T., Duarte, A.C., 2015. A critical overview of the analytical approaches to the occurrence, the fate and the behavior of microplastics in the environment. TrAC - Trends Anal. Chem. 65, 47–53. https://doi.org/10.1016/j.trac.2014.10.011

Rodríguez-Romeu, O., Constenla, M., Carrassón, M., Campoy-Quiles, M., Soler-Membrives, A., 2020. Are anthropogenic fibers a real problem for red mullets (*Mullus barbatus*) from the NW Mediterranean? Sci. Total Environ. 733, 139336. https://doi.org/10.1016/j.scitotenv.2020.139336

Savoca, S., Bottari, T., Fazio, E., Bonsignore, M., Mancuso, M., Luna, G. M.,… & Spanò, N. 2020. Plastics occurrence in juveniles of *Engraulis encrasicolus* and *Sardina pilchardus* in the Southern Tyrrhenian Sea. Science of the Total Environment, 718, 137457. https://doi.org/10.1016/j.scitotenv.2020.137457

Schrank, I., Trotter, B., Dummert, J., Scholz-Böttcher, B.M., Löder, M.G.J., Laforsch, C., 2019. Effects of microplastic particles and leaching additive on the life history and morphology of Daphnia magna. Environ. Pollut. 255. https://doi.org/10.1016/j.envpol.2019.113233

Teles, M., Balasch, J.C., Oliveira, M., Sardans, J., Peñuelas, J., 2020. Insights into nanoplastics effects on human health. Sci. Bull. 65, 1966–1969. https://doi.org/10.1016/j.scib.2020.08.003

Teuten, E.L., Saquing, J.M., Knappe, D.R.U., Barlaz, M.A., Jonsson, S., Björn, A., Rowland, S.J., Thompson, R.C., Galloway, T.S., Yamashita, R., Ochi, D., Watanuki, Y., Moore, C., Viet, P.H., Tana, T.S., Prudente, M., Boonyatumanond, R., Zakaria, M.P., Akkhavong, K., Ogata, Y., Hirai, H., Iwasa, S., Mizukawa, K., Hagino, Y., Imamura, A., Saha, M., Takada, H., 2009. Transport and release of chemicals from plastics to the environment and to wildlife. Philos. Trans. R. Soc. B Biol. Sci. 364, 2027–2045. https://doi.org/10.1098/rstb.2008.0284

Ugwu, K., Herrera, A., Gómez, M., 2021. Microplastics in marine biota: A review. Mar. Pollut. Bull. 169, 112540. https://doi.org/10.1016/j.marpolbul.2021.112540

Wesch, C., Bredimus, K., Paulus, M., Klein, R., 2016. Towards the suitable monitoring of ingestion of microplastics by marine biota: A review. Environ. Pollut. 218, 1200–1208. https://doi.org/10.1016/j.envpol.2016.08.076

Woodall, L.C., Sanchez-Vidal, A., Canals, M., Paterson, G.L.J., Coppock, R., Sleight, V., Calafat, A., Rogers, A.D., Narayanaswamy, B.E., Thompson, R.C., 2021. The deep sea is a major sink for microplastic debris. R. Soc. Open Sci. 1, 140317. https://doi.org/10.1098/rsos.140317

Zhang, L., Li, X., Zhang, S. et al. 2021. Micro-FTIR combined with curve fitting method to study cellulose crystallinity of developing cotton fibers. Anal Bioanal Chem 413, 1313–1320. https://doi.org/10.1007/s00216-020-03094-6.

Zitouni, N., Bousserrhine, N., Belbekhouche, S., Missawi, O., Alphonse, V., Boughatass, I., Banni, M., 2020. First report on the presence of small microplastics (≤ 3 μm) in tissue of the commercial fish Serranus scriba (Linnaeus. 1758) from Tunisian coasts and associated cellular alterations. Environ. Pollut. 263, 114576. https://doi.org/10.1016/j.envpol.2020.114576

