## Supplementary material for "Presence and the global implications of plastics in wild commercial fish in the Alboran Sea": supp table

| Table 1. Specific details of studies assessing plastic ingestion in fishes from 2013 to 2021. | | | | | | | | | | |
| --- | --- | --- | --- | --- | --- | --- | --- | --- | --- | --- |
| Reference | Group | Subgroup | Species | Number of specimens/% with plastics | Location | Sample processing | Analysis technique | Type/size/color of plastics | Concentration (average per ind.) | Contamination control (blanks) |
| *Scyliorhinus canicula* | | | | | | | | |  |  |
| Alomar et al., 2020 | Fish | Demersal fish | *Scyliorinhinus canicula* | 13/8 | Western Mediterranean Sea | gastrointestinal tracts | Stereomicroscope | N.A. | 0.08 | Yes |
| Bellas et al., 2016 | Fish | Demersal fish | *Scyliorinhinus canicula* | 72/15.3 | Spanish Atlantic and Mediterranean coasts | Stomach content | Stereomicroscope | Microplastics (100%), Black (51%) > Red (13%) > Gray (12.70%) > Blue (8.70%) > Brown (6.30%) > Green (4.20%) > Pink (2.10%) > Yellow (2%), 0.38 – 3.1 mm | 1.13 | No |
| Capillo et al., 2020 | Fish | Demersal fish | *Scyliorinhinus canicula* | 12/33 | Southern Tyrrhenian Sea | Digestive tract and gills | Stereomicroscope, MicroRaman | Macroplastics (10%) < Microplastics (90%), Fibers (100%), Black (80%) > Red (20%), Others (50%) > PP (31.25%) > PA (12.5%) > PE (6.25%) | 1.1 | Yes |
| Mancia et al., 2020 | Fish | Demersal fish | *Scyliorinhinus canicula* | 50/80 | Southern Sicilia | digestive tract | Stereomicroscope and Raman | Macroplastic (8%) < Microplastics (92%), Fibers (85.7%) > Fragments (16.7%), Dark color (63.8%) > Light color (21.7%) > Transparent (14.5%), 0.001-0.1 mm | 1.18 | Yes |
| Morgan et al., 2021 | Fish | Demersal fish | *Scyliorinhinus canicula* | 200/13 | West coast (UK) | Stomach and spiral valve | Stereomicroscope | Macroplastic (78.5%) > Microplastic (21.5%), Gray (46%) > Orange (21%) > White (14%) = Brown (14%) > Green (5%) | N.A | No |
| Neves et al., 2015 | Fish | Demersal fish | *Scyliorinhinus canicula* | 20/20 | Coasts of Portugal | Stomach Content | µFTIR | Fibers (83%) > Fragments (17%). 0.217 - 4.81 mm | N.A | No |
| Parton et al., 2020 | Fish | Demersal fish | *Scyliorinhinus canicula* | 12/66.6 | North-East Atlantic (UK) | Digestive tract and stomach content | Stereomicroscope and FTIR | N.A | N.A | Yes |
| Péda et al., 2020 | Fish | Demersal fish | *Scyliorinhinus canicula* | 27/22.2 | Tyrrhenian Sea (Italy) | digestive tract | Stereomicroscope and FTIR | Macroplastic (36%) > Microplastic (64%), Foam (40%) > Fibers, Fragments, Others (20%), White (60%) > Brown (20%) = transparent (20%), PS (40%) > PP (20%) = PA (20%) = PE (20%) | 0.19 | Yes |
| Smith, L. E., 2018 | Fish | Demersal fish | *Scyliorinhinus canicula* | 20/15 | North Sea | Stomach and spiral valve | Stereomicroscope | Macroplastic (66%) > Microplastic (33%), Fibers (33%) = Beads (33%) = Fragments (33%), 1.5 mm | 0.15 | No |
| Study site. 2021 | Fish | Demersal fish | *Scyliorinhinus canicula* | 51/9.8 | Alboran Sea (NW) | Digestive tract | Digestion, Stereomicroscope and FTIR | Macroplastics (60%) > Microplastics (40%), Fibers (100%), Black = Blue (42.86%) > Orange (14.29%), Cellulose (60%) > PVC = PU (20%), 1.037 – 11.18 mm | 0.3 | Yes |
| Valente et al., 2019 | Fish | Demersal fish | *Scyliorinhinus canicula* | 30/66.7 | Tyrrhenian Sea (Italy) | Digestive tract | FTIR | Fibers (85.7%) > Fragments (9.3) > Films (3.5%) > Pellets (1.6%), Dark Color (69.20%) > Light color (22.80%) | 2,5 | Yes |
| Valente et al., 2020 | Fish | Demersal fish | *Scyliorinhinus canicula* | 33/3 | Tyrrhenian coast of central Italy | Digestive tract | Stereomicroscope, and ATR-FTIR | Films (100%), PP (100%) | 0.33 | No |
| *Mullus barbatus* | | | | |  |  |  |  |  |  |
| Anastasopoulou et al. 2018 | Fish | Demersal fish | *Mullus barbatus* | 48/0 | North Adriatic Sea | Stomach and intestine content | Stereomicroscope |  | N.A | Yes |
| Anastasopoulou et al. 2018 | Fish | Demersal fish | *Mullus barbatus* | 50/14 | South Adriatic Sea | Stomach and intestine content | Stereomicroscope |  | N.A | Yes |
| Anastasopoulou et al. 2018 | Fish | Demersal fish | *Mullus barbatus* | 75/16 | NE Ionian Sea | Stomach and intestine content | Stereomicroscope | Macroplastics (33%) < Microplastics (67%), Fibers (78%) > Films (11%) = Beads (11%) | N.A | Yes |
| Avio et al. 2020 | Fish | Demersal fish | *Mullus barbatus* | 10/60 | Nothern Adriatic sea | Digestive tract | μFTIR |  | 2.67 | Yes |
| Avio et al. 2020 | Fish | Demersal fish | *Mullus barbatus* | 10/20 | Central Adriatic sea | Digestive tract | μFTIR |  | 4.37 | Yes |
| Avio et al. 2020 | Fish | Demersal fish | *Mullus barbatus* | 8/37.5 | Southern Adriatic sea | Digestive tract | μFTIR |  | 5.37 | No |
| Bellas et al. 2016 | Fish | Demersal fish | *Mullus barbatus* | 128/18.8 | Spanish Atlantic and Mediterranean coasts | Stomach content | Stereoscopic microscope | Microplastics (100%), Black (51%) > Red (13%) > Gray (12.70%) > Blue (8.70%) > Brown (6.30%) > Green (4.20%) > Pink (2.10%) > Yellow (2%) 0.38 – 3.1 mm | 1.9 | Yes |
| Capillo et al. 2020 | Fish | Demersal fish | *Mullus barbatus* | 21/46.7 | Southern Tyrrhenian Sea | digestive tract and gills | μRaman and ATR-FTIR | Microplastics (100%), Fibers (100%), Black (100%), PP (31.25%) = Others (31.25) > PA (18.25 %) > PE (6.25) | 0.3 | Yes |
| Digka et al. 2018 | Fish | Demersal fish | *Mullus barbatus* | 25/32 | Northern ionian sea | Digestive tract | Stereomicroscope, FT-IR | Fragments (83.3%) > Fibers (17.7), Blue (50%) > Pink (40%) > Green (10%), PE (30%) > PP (20%) = Others (20%) > PS (10%) = PET (10%) 0.048 – 0.8 mm | 1.5 | Yes |
| Giani et al. 2019 | Fish | Demersal fish | *Mullus barbatus* | 132/19.7 | Eastern Sicily | Digestive tract | Stereomicroscope | Microplastics (66%) > Macroplastics (19%), Fibres (44%) > Fragments (32%) > Films (24%), Blue (36%) > Black (25%) > White (21%) > Transparent (14%) > Red (4%) | 1.08 | Yes |
| Gündoğdu et al. 2020 | Fish | Demersal fish | *Mullus barbatus* | 63/63 | Turkish coast | Digestive tract | Stereomicroscope, Raman | Microplastics (100%), Fibers (53.6%) > Fragments (46.4%), Others (41.2%) > PP (26%) > PE (21.9%) > PET (8.2%) > PVC (1.4%) = PA (1.4%), 0.028 – 4.909 mm | 1.1 | Yes |
| Palazzo et al. 2021 | Fish | Demersal fish | *Mullus barbatus* | 89/22.47 | Western Mediterranean Sea | Digestive tract | stereomicroscope and FTIR | Fibers (66.53%) > Fragments (27.31%) > Films (3.46%) > Foam (1.54%) > Beads (1.15%), Blue (37%) > Black (31%) > Transparent (14%) > Green (9%) > Green (3%) > Clear = White = Red (2%), 0.102 – 14.742 mm | 1.8 | Yes |
| Rodríguez-Romeu et al. 2020 | Fish | Demersal fish | *Mullus barbatus* | 118/50 | North-west mediterranean sea | Stomach content | Visual obs. Raman | Microplastics (100%), Fibers (96%) > Fragments (4%), Others (56.79%) > PET (31.14%), 0.37 – 14.80 | 1.48 | No |
| Study site. 2021 | Fish | Demersal fish | *Mullus barbatus* | 52/32.69 | Alboran Sea (NW) | Digestive content | Digestion, Stereomicroscope and FTIR | Microplastics (100%), Fibers (100%), Black (66.67%) > Blue (12.5%) > Gray (8.33%) > Brown = Multicolor = Green (4.17%), Cellulose (66.6%) > Neoprene = PET (16.6%), 1.128 – 8.014 | 0.46 | Yes |
| *Scomber scombrus* | | | | | | | | | | |
| Akoueson et al. 2020 | Fish | Pelagic fish | *Scomber scombrus* | 10/NA | NA | Gills, Guts and flesh tissue | visual obs., micro-FT-IR | Fibers (90%) > Fragments (7%) > Films (3%), Others (58%) > PET (25%) > PP (9%) > PA (8%) | NA | No |
| Avio et al. 2020 | Fish | Pelagic fish | *Scomber scombrus* | 10/90 | Northern Adriatic Sea | Digestive tract | Density separation, Stereomicroscope, μFTIR | Microplastics (100%), Fibers (75%) > Others (25%), | 4.22 | Yes |
| Avio et al. 2020 | Fish | Pelagic fish | *Scomber scombrus* | 10/70 | Central Adriatic sea | Digestive tract | Density separation, Stereomicroscope, μFTIR | Microplastics (100%), Others (shape) (100%) | 1,3 | Yes |
| Foekema et al. 2013 | Fish | Pelagic fish | *Scomber scombrus* | 84/0 | North sea | Digestive tract content | stereomicroscope |  | NA | No |
| Nelms et al. 2018 | Fish | Pelagic fish | *Scomber scombrus* | 31/32 | Celtic sea | Digestive tract | Visual observation, FTIR | Microplastics (100%), Fibers (72%) > Fragments (28%), Blue = Red (28%) > Black (22%) > Orange = Green (11%), Others (60%) > PE (28%) > PA = PP (6%), 0.5 - 0.6 mm | 0.58 | Yes |
| Neves et al. 2015 | Fish | Pelagic fish | *Scomber scombrus* | 13/31 | Coast of Portugal | Stomach content | μFTIR | Microplastics (100%), Fibers (50%) = Fragments (50%), 0.217 – 4.81 mm | 0.46 | No |
| Palazzo et al. 2021 | Fish | Pelagic fish | *Scomber scombrus* | 65/49.23 | Western Mediterranean Sea | Digestive tract | stereomicroscope and FTIR | Fibers (58.9%) > Fragments (30.82%) > Films (6.16%) > Foam (2.71%) > Beads (1.37%), Blue (27%) > Black (24%) > transparent (19%) > Green (8%) > Clear(6%) > Orange = White (5%) > Pink (3%) > Purple = Red = Multicolor (1%), PE (42.11%) > PP = Others (26.32%) > PS (5.26), 0.345 – 19 mm | 2.16 | Yes |
| Rummel et al. 2016 | Fish | Pelagic fish | *Scomber scombrus* | 38/13.2 | North sea | Digestive tract | Density separation, Stereomicroscope, ATR-FTIR | Microplastics (66.7) > Macroplastics (33.3), Fragments (44.2%) > Fibers = Films (22.2) > Beads (11.1%). White and clear (66.6%) > Blue (22.2%) > Black (11.1%), PE (33.3%) > PP = PS = PA (22.2%), 1.66 mm | 1.8 | Yes |
| Rummel et al. 2016 | Fish | Pelagic fish | *Scomber scombrus* | 13/30.8 | Baltic sea | Digestive tract | Density separation, Stereomicroscope, ATR-FTIR | Microplastics (71.43%) > Macroplastics (28.57%), Fragments (71.43%) > Fibers = Films (14.29%), Clear (42.26%) > Green = Brown = Blue = Black (14.29%), PE (57.14%) > PA (28.57%) > PP (14.29%), 2.05 mm | 1.75 | Yes |
| Study site, 2021 | Fish | Pelagic fish | *Scomber scombrus* | 40/35 | Almeria’s Bay | Digestive Tract | Digestion, Stereomicroscope, FTIR | Microplastics (80%) > Macroplastics (20%), Fibers (98%) > Fragments (2%), White and clear (25.7%) > Blue (25.1%) > Black (23.5%) > Green (7.7%) > Red (5.3%) > Orange (5.1%) > Brown (3.3%) > (2.4%) > Others (1.9), Cellulose (60%) > PVC = PE = PET = PP (10%), 1.151 – 21.747 mm | 0.37 | Yes |
| *Engraulis encrasicolus* | | | | | | | | |  |  |
| Alomar et al. 2020 | Fish | Pelagic fish | *Engraulis encrasicolus* | 24/0 | Western mediterranean sea | Gastrointestinal tract | visual |  | NA | Yes |
| Bakir et al. 2020 | Fish | Pelagic fish | *Engraulis encrasicolus* | 178/57 | South african coastline | Gatrointestinal tract | Stereomicroscope, ATR FTIR | Microplastics (100%) | 1.13 | No |
| Collard et al. 2017 (a) | Fish | Pelagic fish | *Engraulis encrasicolus* | 20/40 | Mediterranean sea | Stomatch | Filtration and Raman | Fragments (52%) > Fibers (48%), Blue (46%) > Transparent (22%) > Black (12%) > Red = White (8%) > Pink (4%), PE (37%) > PP (26%) > PET (16%) > Others (11%) > PS = PA (5%) | NA | Yes |
| Collard et al. 2017 (b) | Fish | Pelagic fish | *Engraulis encrasicolus* | 10/80 | Gulf of lions | Stomatch | Filtration and Raman |  | NA | Yes |
| Compa et al. 2018 | Fish | Pelagic fish | *Engraulis encrasicolus* | 105/14.28 | NW mediterranean to gibraltar | Gastrointestinal tract | Stereomicroscope, μFTIR | Microplastics (100%), Fibers (83%) > Fragments (17%), Blue (46%) > Transparent (22%) > Black (12%) > Red = White (8%) > Pink (4%), Others (50%) > PET (30%) > PE (20%) | 0.18 | No |
| Filgueiras et al. 2020 | Fish | Pelagic fish | *Engraulis encrasicolus* | 15/87 | NW Iberian continental shelf | Stomatch content | Stereomicroscope, μRaman | Microplastics (100%), Fibers (64%) > fragments (36%), Transparent (56%) > Blue (12%) > Red (6%) > Black = Green = White (4%) | 1.92 | Yes |
| Kazour et al. 2019 | Fish | Pelagic fish | *Engraulis encrasicolus* | 10/83.3 | Eastern mediterranean basin (Tripoli) | Gatrointestinal tract | Stereomicroscope, μRaman | Fragments (52%) > Fibers (44%) > Films (4%), PP (25%) = PS (25%) = PA (25%) = PU (25%) | 4.3 | Yes |
| Kazour et al. 2019 | Fish | Pelagic fish | *Engraulis encrasicolus* | 10/83.3 | Eastern mediterranean basin (Beirut) | Gatrointestinal tract | Stereomicroscope, μRaman | Fragments (52%) > Fibers (44%) > Films (4%), PS (45%) > PE (20%) > PET (15%) > PU = Others (10%) | 4.3 | Yes |
| Kazour et al. 2019 | Fish | Pelagic fish | *Engraulis encrasicolus* | 10/83.3 | Eastern mediterranean basin (Sidon) | Gatrointestinal tract | Stereomicroscope, μRaman | Fragments (52%) > Fibers (44%) > Films (4%), PS (45%) > PP(15%) > PU = PE = PET = Others (10%) | 4.3 | Yes |
| Lefebvre et al. 2019 | Fish | Pelagic fish | *Engraulis encrasicolus* | 84/11 | Gulf of lions | Digestive tract | Stereomicroscope, μFTIR | Fibers (99.1%) > Fragments (0.9%), Light color (45%) > Dark color = Blue (22%) > Others (11%), PET (89%) > PE (11%), 0.5 – 1.5 mm | 0.11 | Yes |
| Pennino et al. 2020 | Fish | Pelagic fish | *Engraulis encrasicolus* | 103/60 | Northwestern Mediterranean coast | Stomach content | stereomicroscope | Microplastics (100%) | 1.5 | No |
| Renzi et al. 2019 | Fish | Pelagic fish | *Engraulis encrasicolus* | 80/91 | Adriatic sea | Stomach | Stereomicroscope, ATR FTIR | Microplastics (100%) | NA | No |
| Rios-Fuster et al. 2019 | Fish | Pelagic fish | *Engraulis encrasicolus* | 24/0 | Western mediterranean sea (Balearic islands) | Gatrointestinal tract | Stereomicroscope, no digestion |  | 0 | Yes |
| Rios-Fuster et al. 2020 | Fish | Pelagic fish | *Engraulis encrasicolus* | 15/6.67 | Western mediterranean sea (Peninsular coast) | Gatrointestinal tract | Stereomicroscope, no digestion |  | 0.07 | Yes |
| Savoca et al. 2020 | Fish | Pelagic fish | *Engraulis encrasicolus* | 55/47 | Southern Tyrrhenian Sea | External and complet digestion | Density separation, Stereomicroscope, ATR-FTIR and MicroRaman | Microplastics (100%), Fibers (71.43%) > Fragments (28.57%), Black = Blue (42.85%) > Yellow (14.25%) | 0.26 | Yes |
| Study site. 2021 | Fish | Pelagic fish | *Engraulis encrasicolus* | 51/0 | Almeria’s bay | Digestive tract | Digestion, Stereomicroscope |  | 0 | Yes |
