## Supplementary material for "Presence and the global implications of plastics in wild commercial fish in the Alboran Sea": supp figures

Supplementary information

Table 1

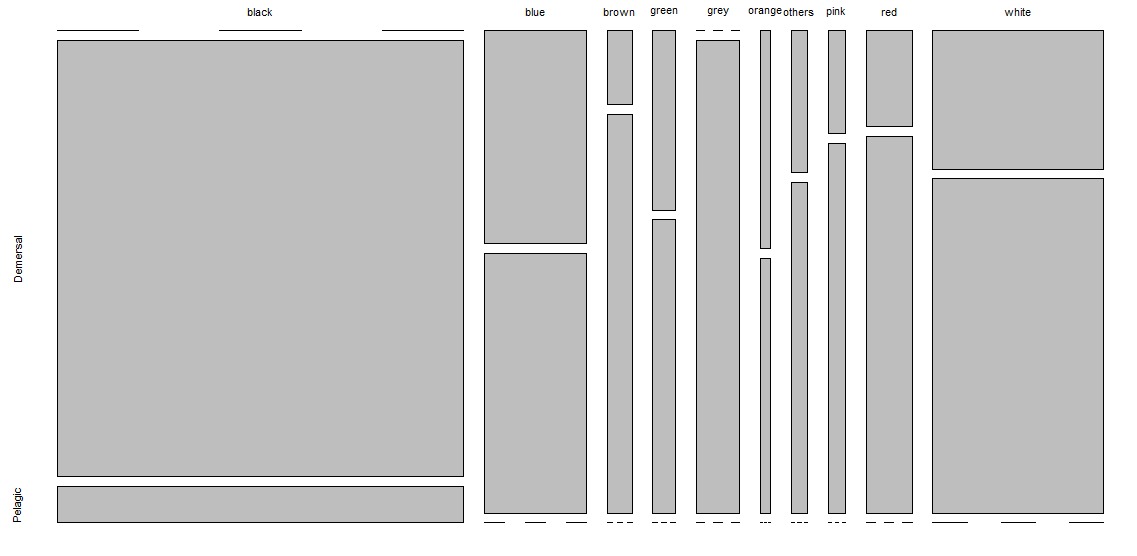

Fig 1. Color frequency among pelagic and demersal group.

a

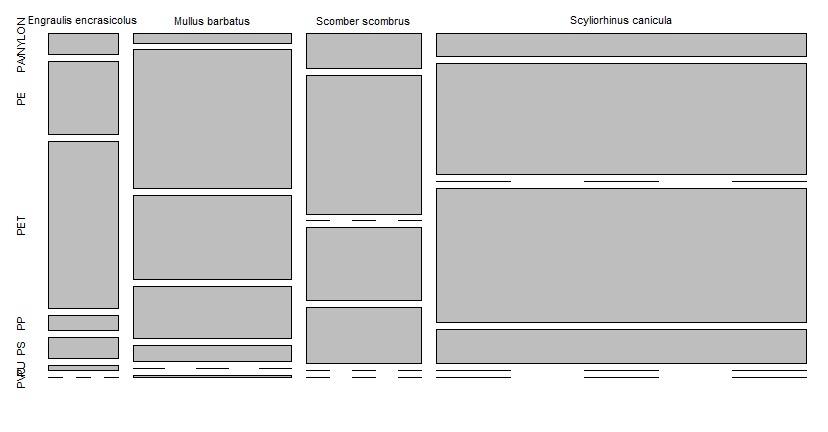

b

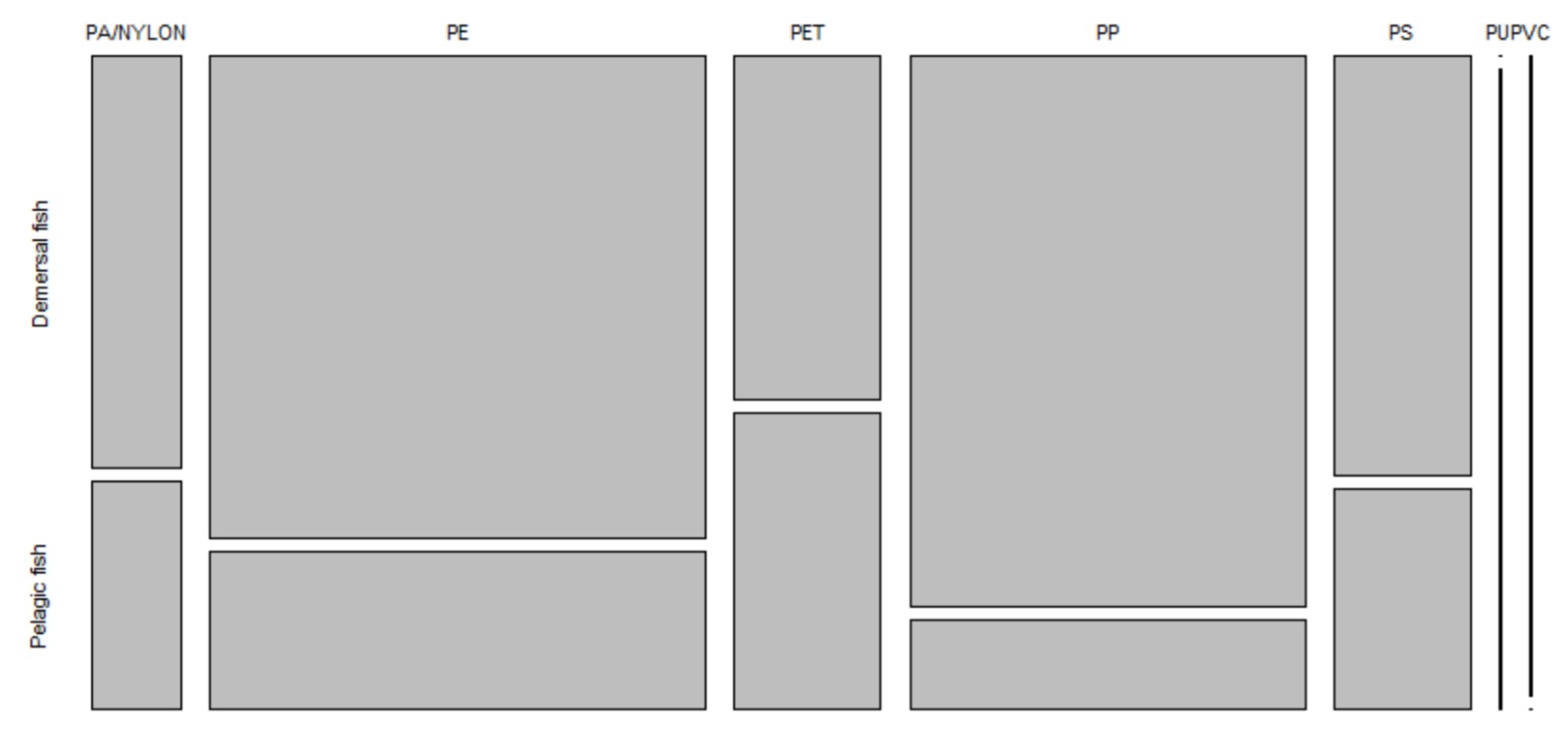

Fig 2. (a) Polymer prevalence by species and (b) between groups
